# Separable mechanisms drive local and global polarity establishment in the *C. elegans* intestinal epithelium

**DOI:** 10.1101/2021.11.01.466827

**Authors:** Melissa A. Pickett, Maria D. Sallee, Victor F. Naturale, Deniz Akpinaroglu, Joo Lee, Kang Shen, Jessica L. Feldman

**Affiliations:** Department of Biology, Stanford University, Stanford, CA 94305, USA

**Keywords:** apico-basolateral polarity, Par3/PAR-3, aPkc/PKC-3, intestine, *C. elegans*

## Abstract

Apico-basolateral polarization is essential for epithelial cells to function as selective barriers and transporters, and to provide mechanical resiliency to organs. Epithelial polarity is established locally, within individual cells to establish distinct apical, junctional, and basolateral domains, and globally, within a tissue where cells coordinately orient their apico-basolateral axes. Using live imaging of endogenously tagged proteins and tissue specific protein depletion in the *C. elegans* embryonic intestine, we found that local and global polarity establishment are temporally and genetically separable. Local polarity is initiated prior to global polarity and is robust to perturbation. PAR-3 is required for global polarization across the intestine but is not required for local polarity establishment as small groups of cells are able to correctly establish polarized domains in PAR-3 depleted intestines in an HMR-1/E-cadherin dependent manner. Despite belonging to the same apical protein complex, we additionally find that PAR-3 and PKC-3/aPKC have distinct roles in the establishment and maintenance of local and global polarity. Together, our results indicate that different mechanisms are required for local and global polarity establishment *in vivo*.

**SUMMARY STATEMENT:** Live-imaging and intestine specific protein depletion reveal that apico-basolateral polarity establishment can be temporally and genetically separated at the local level of individual cells and globally, across a tissue.

## INTRODUCTION

Epithelial cells form adherent sheets that line metazoan organs where they separate internal and external compartments, act as barriers, selectively transport molecules, and provide mechanical resiliency. Underlying these functions is the characteristic polarization of epithelial cells along an apico-basolateral axis. The apical surface faces the external compartment or hollow lumen and is highly specialized by tissue type. The basolateral domains provide contact between neighboring cells and attachment to underlying basement membranes. Adherens junctions (AJs) and occluding junctions (septate junctions (SJs) in invertebrates and tight junctions in vertebrates) are positioned between the apical and basolateral domains, where they provide cell-cell adhesion and barrier functions.

As epithelia are composed of individual polarized cells that collectively form a functional tissue, polarity establishment occurs both locally, with each individual cell establishing an apico-basolateral axis, and globally, such that neighboring cells coordinately orient their axes in the same direction. Polarity establishment at both levels is critical for organ function. Loss of local cell polarity promotes cell growth in pre-invasive breast carcinomas (Halaoui et al., 2017), even though other cells in the tissue retain correct polarization. Partial or complete inversion of local polarity is associated with microvillus inclusion disease (Michaux et al., 2016), and pathogenic bacteria can appropriate local apico-basolateral polarity programs in the intestine, disrupting barrier function and providing bacteria access to internal compartments (Hua et al., 2018; Tapia et al., 2017a; Tapia et al., 2017b). Loss of global, tissue-level polarity results in non-adherent cells that disassociate or fail to undergo morphogenic movements, resulting in embryonic lethality (Achilleos et al., 2010; Bilder et al., 2000; Bossinger et al., 2001; Hutterer et al., 2004; Legouis et al., 2000; McMahon et al., 2001; Totong et al., 2007) or that promote the progression and metastasis of cancers (Catterall et al., 2020; Ellenbroek et al., 2012; Halaoui et al., 2017; Zhou et al., 2017).

Despite the conservation of polarity proteins across species and epithelial tissues, the pathways by which these proteins are deployed and their requirement for polarity establishment are surprisingly divergent (Pickett et al., 2019). The best understood polarity programs have canonically been regarded as “top-down” pathways, where the apical scaffolding protein, Par3/PAR-3/Baz, is the most upstream known polarity player and is necessary to establish apical, junctional, and basolateral domains (Achilleos et al., 2010; Harris and Peifer, 2004; Harris and Peifer, 2005). In these programs, junctional proteins are required for refining polarity and basolateral proteins play downstream roles in polarity maintenance (Bilder et al., 2000; Bilder et al., 2003; Harris and Peifer, 2004; Legouis et al., 2000; McMahon et al., 2001).

However, holistic examination of polarity proteins at different stages in *Drosophila* blastoderm development have revealed that such “top-down” pathways are more complex and less linear than originally thought. Par3 acts both upstream and downstream of the AJ protein Afadin/Canoe (Bonello et al., 2018). Similarly, the basolateral scaffolding proteins Dlg and Scribble were found to play essential roles in the initial positioning of AJs (Bonello et al., 2019). Additionally, other epithelia can polarize independently of conserved polarity proteins or through parallel, redundant mechanisms as in the *Drosophila* midgut and follicular epithelium, respectively (Chen et al., 2018; Fernandez-Minan et al., 2008; Pickett et al., 2019; Schneider et al., 2006; Tanentzapf et al., 2000). Thus, closer examination of how different epithelia polarize *in vivo* will provide a more complete understanding of polarity establishment pathways. As ubiquitous removal of polarity proteins from embryonic tissues often results in pleiotropic defects and embryonic lethality, tissue specific protein depletion is critical for understanding the range and requirement of different polarity programs for epithelial tissue structure and function.

To understand the interplay of polarity proteins during local and global apico-basolateral polarity establishment, we characterized the localization of polarity proteins relative to one another and degraded individual polarity proteins specifically from the developing embryonic *C. elegans* intestine. The *C. elegans* intestine is a simple epithelium derived from a single blastomere (“E”) that undergoes four rounds of division, giving rise to a 16-cell intestinal primordium (“E16”) (Leung et al., 1999). At this E16 stage, 10 dorsally positioned and 6 ventrally positioned cells surround a central midline, the future site of the apical surface of each cell and of the intestinal lumen (Leung et al., 1999). Proteins bound for the apical surface first concentrate locally as puncta on lateral membranes and then spread along the central midline, thereby beginning the specification of the global polarity axis in the intestine (Achilleos et al., 2010; Feldman and Priess, 2012; Totong et al., 2007). PAR-3 is the most upstream known polarity protein in the intestine, required for the apical localization of the conserved PAR complex proteins PAR-6 and PKC-3/aPKC and for positioning of junctional proteins (Achilleos et al., 2010). Junctional and some basolateral proteins appear to be required for the maintenance, but not establishment, of polarity, but when and how these proteins become organized into distinct domains is largely uncharacterized (Bossinger et al., 2004; Legouis et al., 2000; McMahon et al., 2001). Furthermore, whether local and global polarity depend on shared or separable mechanisms has not been investigated.

We found that local and global polarity establishment are separable processes in the *C. elegans* embryonic intestine. Local polarity establishment began prior to global establishment, with asymmetric localization of apical and AJ proteins into lateral puncta. In control embryos, global polarity appeared to arise in a stepwise manner with the apical surface established first, followed by the ladder-like junctional structures, and lastly the basolateral domain. However, intestine-specific depletion of polarity proteins revealed that polarity establishment is not a simple linear process. We found that PAR-3 and PKC-3/aPkc are both required for the formation of a functional intestine with a continuous, hollow lumen and for larval growth and viability, but that PAR-3 and PKC-3 play different roles in polarization. PKC-3 was not required to establish apico-basolateral polarity at the local or global levels but was required for SJ protein organization and for maintaining apical and junctional continuity. In contrast, PAR-3 was required for the initial establishment of polarity at both the local and global levels. Intriguingly, older PAR-3 depleted embryonic intestines contained small discontinuous regions with the correct relative organization of apical, junctional, and basolateral proteins, suggesting that local polarity was eventually established. These structures were lost when HMR-1/E-cadherin was co-depleted with PAR-3, indicating that HMR-1 and PAR-3 redundantly drive local polarity establishment. Together, these results reveal that PAR-3 is required for global, but not local polarity establishment in the intestine, and that local polarity establishment is robust and separable from global polarity establishment.

## RESULTS

### Apical and basolateral domains are established at different times and through different routes

To systematically characterize the relative localization of polarity proteins over the course of local and global polarity establishment, we live imaged endogenously tagged apical, basolateral, and junctional proteins in the *C. elegans* embryonic intestine. We restricted our analysis from the appearance of local polarity at the beginning of E16 through the comma stage, when apical, junctional, and basolateral proteins are known to be globally polarized (Achilleos et al., 2010; Beatty et al., 2010; Legouis et al., 2000). We found that four embryonic stages provided clear snapshots of local and global polarity establishment which we defined based on embryo morphology and PAR-3 localization: 1) Stage 1, early pre-bean, ∼330 minutes post fertilization (mpf); 2) Stage 2, mid pre-bean, ∼350 mpf; 3) Stage 3, early bean, ∼390 mpf; 4) Stage 4, comma, ∼430 mpf (Figs. 1A-A’’’, S1).

**Figure 1.**
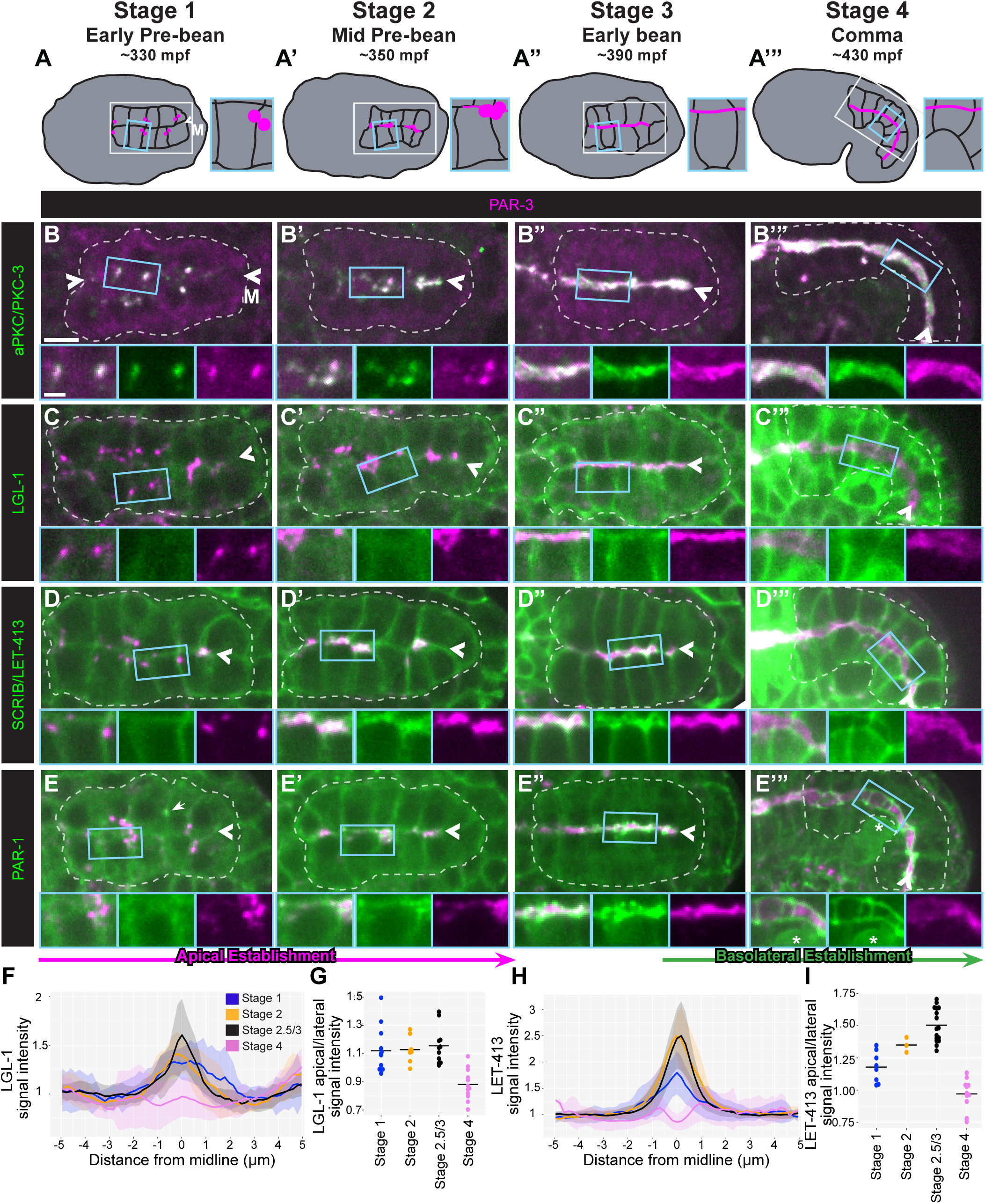
Apical and basolateral proteins localize to the intestinal midline during global polarity establishment. A-A’’’) Schematics of *C. elegans* embryonic morphogenesis from early pre-bean (Stage 1) to comma (Stage 4), with minutes post-fertilization (mpf) indicated. Intestinal membranes in black lines, intestines marked with white boxes, and enlarged insets shown with blue boxes. B-E’’’) Dorsal view of co-localization of endogenously tagged indicated proteins (green) and PAR-3 (magenta) in Stage 1 (PKC-3, n=12; LGL-1, n=12 ; LET-413, n=11; PAR-1, n=10), Stage 2 (PKC-3, n=11; LGL-1, n= 9; LET-413, n=8; PAR-1, n=12), Stage 3 (PKC-3, n=5; LGL-1, n=9 ; LET-413, n=13; PAR-1, n=11) or Stage 4 (PKC-3, n=16; LGL-1, n=12 ; LET-413, n=15; PAR-1, n=12) intestines. Note that PAR-1 occasionally localized to midbodies (E, arrow) in Stage 1 and was highly expressed in germ cells (E’’’, asterisk). All images are maximum intensity projections from live imaging. Intestines outlined by white dashed lines, midlines indicated by arrowheads. Enlarged versions of boxed regions shown below. Scale bar = 5 μm for panels and 2 μm for boxed insets. F-G) Average profile lines for LGL-1 (F) or LET-413 (G) signal across the intestinal midline from Stages 1-4. H-I) Quantification of the apical/lateral LGL-1 (H) or LET-413 (I) signal.

Local polarity was initiated when PAR complex proteins PAR-3 and PKC-3 co-localized in puncta on lateral cell membranes between adjacent non-sister cells (Fig. 1B, Stage 1), consistent with previous observations (Achilleos et al., 2010; Feldman and Priess, 2012). These ‘lateral puncta’ moved toward the intestinal midline, where they spread within each cell (Fig. 1B’, Stage 2), creating a continuous apical surface along the intestinal midline and establishing global apical polarity (Fig. 1B’’, Stage 3) that persisted throughout intestinal morphogenesis (Fig. 1B’’’, Stage 4).

In contrast, basolateral proteins first localized to all plasma membranes, including the apical membrane, before becoming restricted in their localization. LGL-1/Lgl and LET-413/Scribble initially localized weakly to all cell membranes and were neither localized to nor excluded from the lateral puncta (Fig. 1C, D, F-I, Stage 1), becoming more strongly associated with all plasma membranes as polarization progressed (Fig. 1C’,D’, F-I, Stage 2). Surprisingly, LGL-1 and LET-413 colocalized with PAR-3 at the nascent apical surface (Fig. 1C’’,D’’, F-I, Stage 3) before being excluded by Stage 4, as previously described (Fig. 1C’’’, D’’’, F-I) (Beatty et al., 2010; Bossinger et al., 2004; Legouis et al., 2000).

We also examined the localization of the conserved kinase PAR-1/Par1, which localizes to basolateral membranes in many epithelia. We tagged endogenous PAR-1 with GFP and found that it localized to centrosomes during mitosis prior to Stage 1 (Data not shown) and occasionally at midbodies in Stage 1 embryos (Fig. 1E), consistent with the ability of PAR-1 to bind to microtubules and associate with microtubule-binding proteins (Sanchez (Cohen et al., 2004; Doerflinger et al., 2003; Sanchez et al., 2021). At Stage 1, PAR-1 localized weakly to plasma membranes (Fig. 1E), similar to LGL-1 and LET-413. Unlike LGL-1 or LET-413, PAR-1 separated from PAR-3 in Stage 2 (Fig. 1E’). PAR-1 then localized into subapical bands, losing its lateral localization (Fig, 1E’’, Stage 3) and organizing into ladder-like junctions in Stage 4 (Fig. 1E’’’). Thus, PAR-1 shows junctional rather than basolateral localization in the *C. elegans* intestine similar to DLG-1 (Bossinger et al., 2001).

### Adherens and septate junction proteins localize separately during polarity establishment

We next examined the localization of junctional proteins during intestinal polarization. In the *C. elegans* intestine, AJ and SJ proteins appear to occupy separate domains of a single electron-dense junctional structure, the “apical junction” (Koppen et al., 2001; Segbert et al., 2004). We therefore tested whether AJ and SJ proteins showed similar or distinct localization patterns during polarity establishment. We found that the AJ protein HMR-1/E-cadherin co-localized with PKC-3 in lateral puncta in Stage 1 intestines and both moved together toward (Fig. 2A) and subsequently spread along the midline (Fig. 2A’, Stage 2). Shortly after spreading across the midline, HMR-1 shifted into bands parallel to the apical surface (Fig. 2A’’, Stage 3) forming the characteristic ladder-like junctions in older embryos (Fig. 2A’’’, Stage 4). We next explored the localization of AFD-1/Afadin/Canoe, a conserved AJ protein that is required for correct localization of Par3/Baz in the *Drosophila* blastoderm (Bonello et al., 2018; Choi et al., 2013), which was uncharacterized in the *C. elegans* intestine. We tagged endogenously expressed AFD-1 with GFP and found that like HMR-1, it localized to lateral puncta that moved toward the midline (Fig. 2 B, B’ Stages 1 and 2) before separating into subapical bands (Fig. 2B’’, Stage 3) and organizing into ladder-like junctions (Fig. 2B’’’, Stage 4), suggesting AFD-1 is an AJ protein in the *C. elegans* intestine.

**Figure 2.**
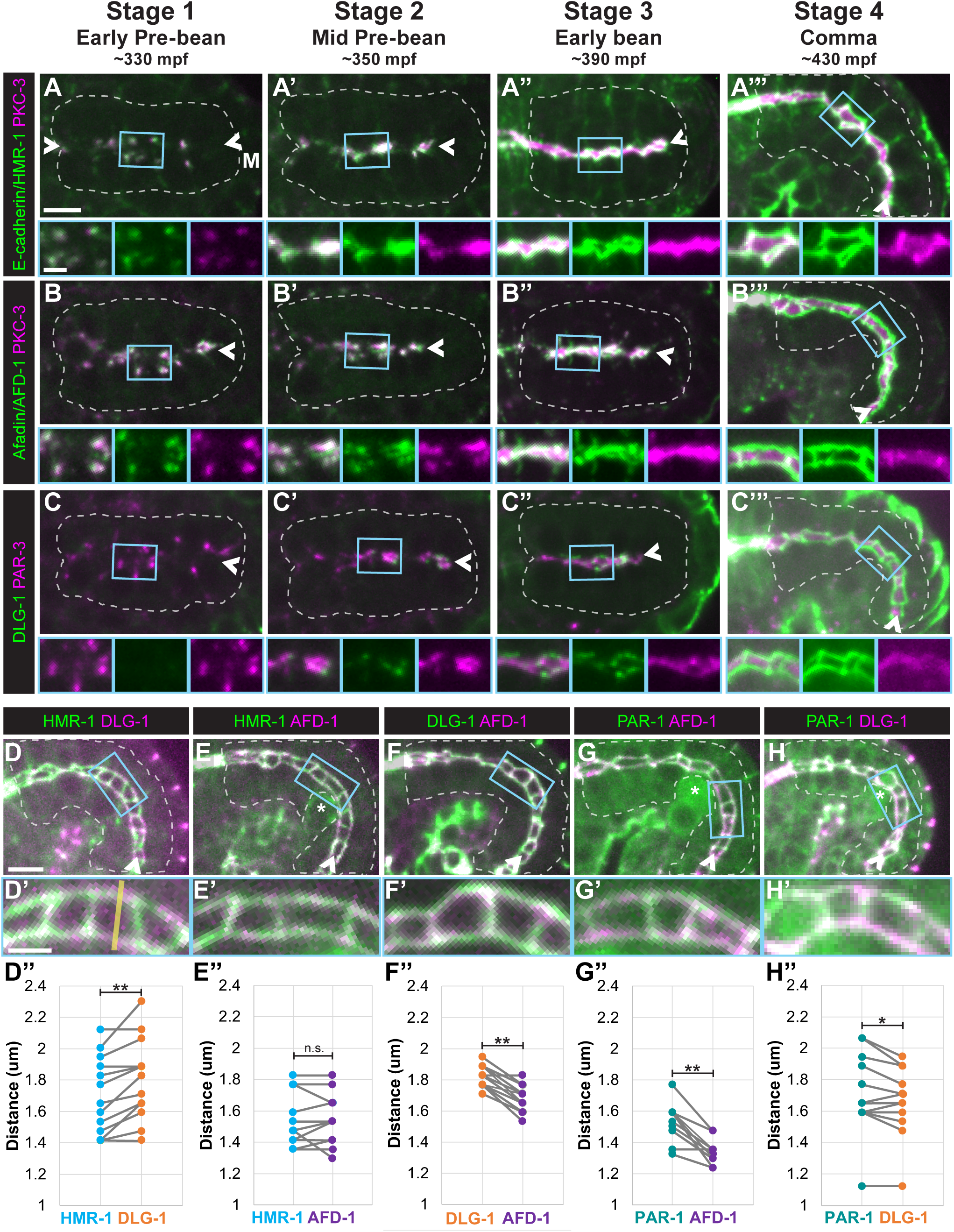
Adherens and Septate-like junctional proteins localize at different times and places during local and global polarity establishment. Dorsal view of co-localization of endogenously tagged indicated proteins (green) and either PKC-3 (A-B, magenta) or PAR-3 (C, magenta) in live Stage 1 (HMR-1, n=11; AFD-1, n=8; DLG-1, n=14), Stage 2 (HMR-1, n=13; AFD-1, n=26; DLG-1, n=13), Stage 3 (HMR-1, n=11; AFD-1, n=8; DLG-1, n=17) or Stage 4 (HMR-1, n=5; AFD-1, n=9; DLG-1, n=12) intestines. D-H) Dorsolateral views of endogenously tagged indicated proteins in live Stage 5 (1.5-fold) intestines (D, n=15; E, n=12; F, n=15; G, n=12; H, n=14). Asterisks mark germ cells. Insets of boxed regions shown below. All images are maximum intensity projections from live imaging. Intestines outlined by white dashed lines, midlines indicated by arrowheads. Scale bar = 5 μm for panels and 2 μm for boxed insets. D’’-H”) Quantification of the distance between the left and right sides of the HMR-1 and DLG-1 junctional structures, measured with a profile line for signal intensity as shown by the yellow line in D’ (two-tailed paired t-test: *: p=0.0.0216, **: p<0.001, n.s.: not significant).

Although the *C. elegans* apical junction appears as a single structure by electron microscopy, the SJ protein DLG-1 localizes more basally than the AJ protein HMR-1 (Koppen et al., 2001; Segbert et al., 2004). Early differences were apparent at the onset of polarity establishment in Stage 1 as the SJ protein DLG-1 was not observed (Fig. 2C), while HMR-1 and AFD-1 were in lateral puncta. In Stage 2 intestines, DLG-1 localized in subapical puncta that were adjacent to, but not co-localized with PAR-3 (Fig. 2C’). DLG-1 then formed fragmented subapical bands adjacent to the apical surface (Fig. 2C’’, Stage 3) that appeared less continuous and more basal than HMR-1 and AFD-1 at the same stage. These bands eventually formed ladder-like junctions (Fig. 2C’’’, Stage 4).

To determine how SJ and AJ proteins are organized within the mature junctions, we compared the localization of junctional proteins in 1.5-fold – 1.7-fold stage embryos. We measured the distance across the midline between intensity peaks, corresponding to the left and right sides of the junction, for two junctional markers, pairing the measurements for each embryo to account for inter-embryo variation in junctional width (see Methods, Fig. S2). Confirming the sensitivity of our method, we measured a significantly smaller distance between intensity peaks of the AJ protein HMR-1 than the SJ protein DLG-1 (Fig. 2D-D”), consistent with HMR-1 localizing apical to DLG-1 (Segbert et al., 2004). The distance between AFD-1 peaks was not significantly different than HMR-1 (Fig. 2E-E’’) but was significantly smaller than DLG-1 (Fig. 2F-F’’), consistent with AFD-1 localizing to the more apical AJ-like region of the apical junction. Surprisingly, the PAR-1 distance was significantly greater than both AFD-1 (Fig. 2G-G”) and DLG-1 (Fig. 2H-H”), suggesting PAR-1 may localize to a SJ-like or more basal junctional structure. Thus, these proteins fall into distinct regions of the apical junction with an basal to apical localization pattern of: PAR-1, DLG-1, HMR-1/AFD-1. The distinct localization and separation of AJ and SJ proteins throughout polarity establishment suggests that the apical junction may be more separable into AJ and SJ-like regions than previously appreciated.

The differences in the localization and movement of polarity proteins into distinct apical, junctional, and basolateral domains indicate that multiple mechanisms likely contribute to local and global polarity establishment in the intestine. Additionally, the separation of junctional structures from the apical surface when basolateral proteins are still co-localized with PAR-3, suggests that global apico-basolateral polarity proceeds temporally in a step-wise manner with the apical surface established first, followed by the junctions, and the basolateral domain established last.

### Different apical and junctional proteins play distinct roles in intestine formation and organismal health

Ubiquitous protein depletion strategies have previously identified key proteins required for polarity establishment or maintenance in the *C. elegans* intestine (Achilleos et al., 2010; Bossinger et al., 2001; Koppen et al., 2001; Legouis et al., 2000; McMahon et al., 2001; Segbert et al., 2004; Totong et al., 2007; Von Stetina and Mango, 2015). However, the essential role of these proteins in many embryonic tissues often resulted in early embryonic lethality when depleted that precluded further functional analysis. Therefore, we depleted polarity proteins specifically from the intestine using the ZF/ZIF-1 system (Armenti et al., 2014; Sallee et al., 2018). Using CRISPR/Cas9, we endogenously tagged polarity proteins with ZF:GFP, allowing ZIF-1-mediated degradation via the ZF1 degron tag (Nance et al., 2003). We expressed ZIF-1 under the *elt-2* promoter for efficient intestine-specific protein depletion (“gut(-)”) beginning in embryos at the “E4” 4-celled intestine stage, well before polarization at Stage 1, and continuing through adulthood. Intestine-specific depletion of either of the apical proteins PAR-3 or PKC-3 (PAR-3^gut(-)^ or PKC-3^gut(-)^, Fig. S3) resulted in 100% developmental arrest at the first larval stage (“L1”, Figs. 3A, S4) as previously described for intestine-specific depletion of PAR-6, the binding partner of PKC-3 (Sallee et al., 2021). Similarly, intestine-specific depletion of HMR-1 caused many larvae to arrest as L1 larvae, though ∼30% of HMR-1^gut(-)^ larvae grew more slowly and developed beyond the first larval stage (Fig. 3A). Larvae depleted of other junctional proteins, AFD-1^gut(-)^ or DLG-1^gut(-)^, also grew slower than control larvae, though unlike HMR-1^gut(-)^, most developed beyond the L1 stage.

**Figure 3.**
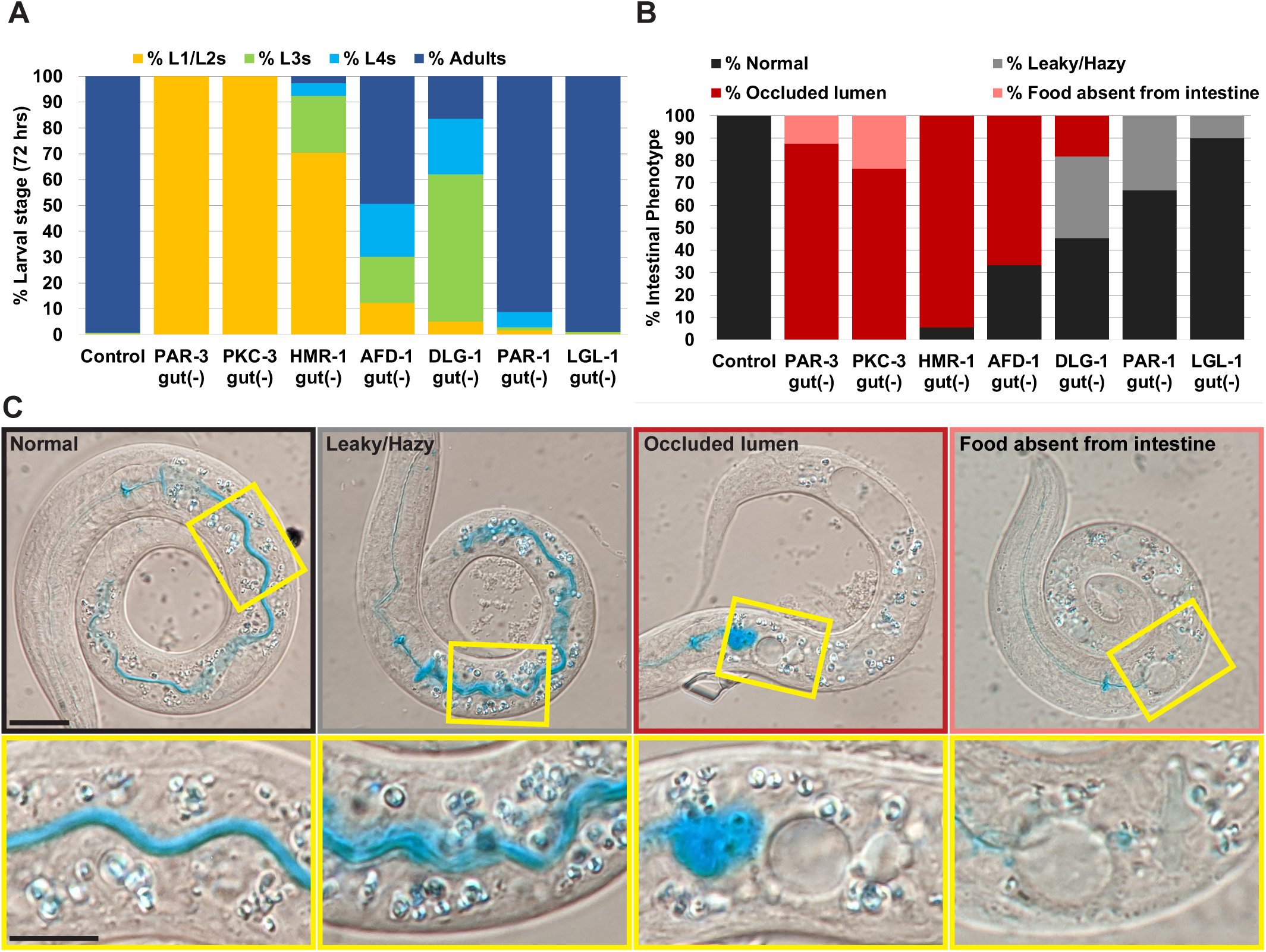
Polarity proteins are differentially required for intestinal structure and function and larval growth. A) Percentage of worms at the L1/L2, L3, or L4 larval or adult stage 72 hours after egg lay for control (*zif-1(gk117);intDeg)* or intestine specific depletion of indicated protein (Control, n=696; PAR-3^gut(-)^, n=198; PKC-3^gut(-)^, n=132; HMR-1^gut(-)^, n=186; AFD-1^gut(-)^, n=146; DLG-1^gut(-)^, n=116; PAR-1^gut(-)^, n=171; LGL-1^gut(-)^, n=269). 100% of PAR-3^gut(-)^ and PKC-3^gut(-)^ worms arrested as L1 larvae. B) Percentage of L1 larvae with normal, leaky/hazy, or occluded intestinal lumens, or entirely lacking food in the intestine for control or following intestine specific depletion of indicated protein (Control, n=13; PAR-3^gut(-)^, n=16; PKC-3^gut(-)^, n=17; HMR-1^gut(-)^, n=18; AFD-1^gut(-)^, n=12; DLG-1^gut(-)^, n=11; PAR-1^gut(-)^, n=15; LGL-1^gut(-)^, n=10). C) Representative DIC images of live worms fed blue food coloring showing indicated phenotypic categories, with higher magnification views of boxed regions shown below. Scale bar = 20 μm for panels and 10 μm for boxed insets.

To determine if intestinal structure and function were compromised following the depletion of apical or junctional proteins, we fed newly hatched L1 larvae bacteria dyed with blue food coloring (“Smurf assay”) which is limited to the continuous, hollow lumen of functional intestines (Fig. 3B, C, (Gelino et al., 2016; Sallee et al., 2021). The majority of PAR-3^gut(-)^ or PKC-3^gut(-)^ larvae showed occluded intestinal lumens, with food pooled posterior to the pharynx and large edemas throughout the intestine (Fig. 3B, C). We occasionally observed small, discontinuous regions resembling lumens in PKC-3^gut(-)^ intestines that we never observed in PAR-3^gut(-)^ larvae (Fig. S4). Most HMR-1^gut(-)^ L1 larvae also had anterior intestinal occlusions (88%) and discontinuous edematous lumens (72%, Figs. 3B,C and S4). Unlike PAR-3^gut(-)^ and PKC-3^gut(-)^ L1 larvae, many HMR-1 L1 larvae had regions of continuous lumens after the anterior occlusion (66%), though these were rarely filled with food. AFD-1^gut(-)^ worms displayed a range of phenotypes, with the majority exhibiting anterior occlusions (58%, Figs. 3B, C, S4), though food filled more of the anterior intestine than in PAR-3^gut(-)^, PKC-3^gut(-)^ or HMR-1^gut(-)^ larvae and the lumens appeared to be continuous posterior to the occlusion (Fig. S4). DLG-1^gut(-)^ larvae also exhibited a range of phenotypes with blue dye often observed outside the lumen in intestinal cells, perhaps indicating barrier dysfunction (36%, Fig. 3B, C Leaky/hazy), but most DLG-1^gut(-)^ larvae had a continuous lumen by DIC and the Smurf assay (75%. Fig. 3B, C). PAR-1^gut(-)^ larvae had still milder defects, with only ∼9% showing a developmental delay (Fig. 3A) and only 33% of PAR-1^gut(-)^ intestines showing possible barrier dysfunction (Figs. 3B and S4). Depletion of the basolateral protein LGL-1 from the intestine did not slow worm growth or perturb intestinal function (Fig. 3A-C), consistent with earlier studies showing that LGL-1 was dispensable for worm growth and survival (Beatty et al., 2010). Together, these results reveal differences in protein requirement to build a functional intestine, with the apical PAR and AJ proteins critical for intestinal function and organismal viability, and SJ and basolateral proteins relatively dispensable.

### Different apical and junctional proteins play different roles in global apical establishment

We next asked how depletion of different apical and junctional proteins affected polarity establishment. We found that PAR-3 was required to localize PKC-3 at all stages of polarity establishment (Fig. S5), consistent with prior studies and indicative of effective PAR-3 degradation (Achilleos et al., 2010). PKC-3 was not required for initial PAR-3 localization (Fig. S5) but was required to maintain continuous PAR-3 localization along the midline, as gaps appeared along this surface in elongating embryos (Fig. S5, (Sallee et al., 2021)). In the *Drosophila* blastoderm, Afadin/Canoe is required for the localization of Par3/Baz. However, consistent with the mild defects observed in AFD-1^gut(-)^ L1 larvae, AFD-1^gut(-)^ embryos had perturbed but relatively normal apical intestinal polarity compared to PAR-3 depleted intestines. Stage 1 AFD-1^gut(-)^ embryos had fewer PAR-3 and PKC-3 lateral puncta, particularly in the anterior part of the intestine where puncta were often absent, and generally reduced fluorescence intensity in the posterior region of the intestine (Fig. S5). This defect in anterior intestinal cells persisted through Stage 2 (Fig. S5), though PKC-3 became correctly localized to the midline beginning in Stage 3. Minor extensions of PKC-3 away from the apical surface were occasionally observed in Stage 4 and older AFD-1^gut(-)^ embryos (Fig. S5), similar to those observed in embryos with actin defects (Gobel et al., 2004; Ramalho et al., 2020; Sanchez et al., 2021; Sasidharan et al., 2018; Van Furden et al., 2004). Importantly, this result suggests that while AFD-1 may contribute to the assembly and movement of lateral puncta to the intestinal midline, other mechanisms are sufficient for local and global apical establishment. By contrast, PAR-3 and PKC-3 lateral puncta formed and moved to the intestinal midline normally in PAR-1^gut(-)^ embryos (Fig. S5). In Stage 4 and older PAR-1^gut(-)^ embryos, we occasionally observed an expansion of the apical surface, consistent with the rare growth defects observed in PAR-1^gut(-)^ larvae (Fig. S5).

### PAR-3 has PKC-3-independent roles in the positioning and maturation of junctions

Given the similarity of the terminal phenotype of PAR-3^gut(-)^ and PKC-3^gut(-)^ larvae, we asked whether their earlier defects in intestinal development were also similar. PAR-6, PAR-3, and PKC-3 homologs are known to play roles in junctional establishment and organization in many systems, so we determined whether PAR-3 regulates junctions through PKC-3 activity or through a separate mechanism by examining the localization of AJ and SJ proteins in PKC-3^gut(-)^ or PAR-3^gut(-)^ embryos (Achilleos et al., 2010; Elbediwy et al., 2019; Iden et al., 2012; Sallee et al., 2021; Totong et al., 2007). AJ proteins HMR-1 and AFD-1 localized near the newly established apical surface along the intestinal midline in both Stage 3 control (Fig. 4A, D) and PKC-3^gut(-)^ embryos (Fig. 4B, E). In Stage 4 control embryos, HMR-1 and AFD-1 localized into ladder-like subapical junctions (Fig. 4A’, A”, D’, D’’), but these junctions appeared to be closer together in Stage 4 PKC-3^gut(-)^ embryos, possibly reflecting a narrower apical surface or inefficient separation of junctions (Fig. 4B’,E’). The AJ phenotypes became more severe in older 1.5-fold PKC-3^gut(-)^ embryos, with more gaps in HMR-1 and AFD-1 observed (Fig. 4B”, E”). This result indicates that PKC-3 is not required to organize AJ proteins but is required for the growth and continuity of the apical surface, like its binding partner PAR-6 (Sallee et al., 2021). By contrast, in Stage 3 PAR-3^gut(-)^ embryos, HMR-1 and AFD-1 localized weakly in patches and along membranes (Fig. 4C, F) instead of as parallel subapical bands. In Stage 4 PAR-3^gut(-)^ intestines, HMR-1 and AFD-1 formed separated rings (Fig. 4C’, F’) that became more continuous and prominent in 1.5-fold intestines (Fig. 4C’’, F’’). Surprisingly, HMR-1 and AFD-1 were excluded from the center of the rings (Fig. 4C”, F”), suggesting that local junction organization may be intact within small cell clusters, and demonstrating that PAR-3 has PKC-3-independent roles in AJ protein localization.

**Figure 4.**
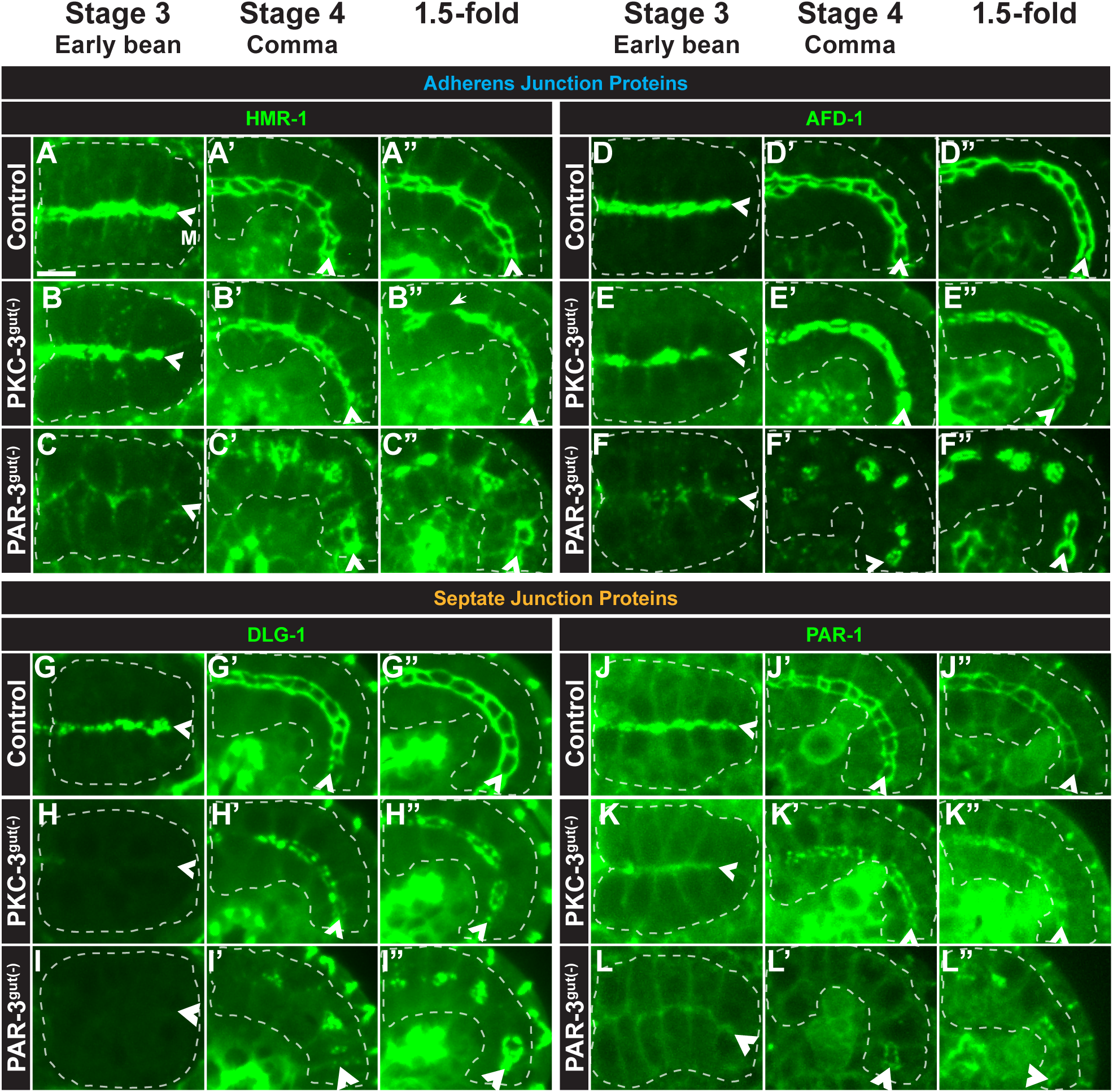
PAR-3 and PKC-3 play different roles in AJ and SJ protein localization. A-L’’) Dorsal (Stage 3) or dorsolateral (Stage 4 and 1.5-fold) live images of indicated endogenously tagged AJ or SJ protein localization in control (A, D, G, J), PKC-3^gut(-)^ (B, E, H, K), or PAR-3^gut(-)^ (C, F, I, L) embryos. B”) Gaps in junctional proteins frequently appeared in 1.5-fold PKC-3^gut(-)^ intestines (B”, arrow). All images are maximum intensity projections from live imaging. Intestines outlined by white dashed lines, midlines indicated by arrowheads. Scale bar = 5 μm. Stage 3 HMR-1 (Control, n=14; PAR-3^gut(-)^, n=10; PKC-3^gut(-)^, n=13); AFD-1 (Control, n=4; PAR-3^gut(-)^, n=3; PKC-3^gut(-)^, n=6); DLG-1 (Control, n=12; PAR-3^gut(-)^, n=1; PKC-3^gut(-)^, n=7); PAR-1 (Control, n=4; PAR-3^gut(-)^, n=2; PKC-3^gut(-)^, n=9). Stage 4 HMR-1 (Control, n=8; PAR-3^gut(-)^, n=18; PKC-3^gut(-)^, n=12); AFD-1 (Control, n=10; PAR-3^gut(-)^, n=7; PKC-3^gut(-)^, n=12); DLG-1 (Control, n=24; PAR-3^gut(-)^, n=12; PKC-3^gut(-)^, n=26); PAR-1 (Control, n=13; PAR-3^gut(-)^, n=16; PKC-3^gut(-)^, n=16). 1.5-fold HMR-1 (Control, n=5; PAR-3^gut(-)^, n=12; PKC-3^gut(-)^, n=13); AFD-1 (Control, n=6; PAR-3^gut(-)^, n=8; PKC-3^gut(-)^, n=10); DLG-1 (Control, n=23; PAR-3^gut(-)^, n=5; PKC-3^gut(-)^, n=19); PAR-1 (Control, n=8; PAR-3^gut(-)^, n=10; PKC-3^gut(-)^, n=8).

We next explored whether SJ proteins were similarly disrupted upon PAR-3 or PKC-3 depletion. In control embryos, PAR-1 and DLG-1 localized in subapical bands in Stage 3 intestines and become localized to the basal part of the junctions in Stage 4 and 1.5-fold intestines (Fig. 4G-G’’, J-J’’). In Stage 3 PKC-3^gut(-)^ intestines, DLG-1 localization was absent, and PAR-1 appeared weakly associated with the apical and lateral cell membranes but was absent from subapical bands (Fig. 4H, K). In Stage 4 PKC-3^gut(-)^ intestines, both DLG-1 and PAR-1 localized weakly and discontinuously into subapical junctions (Fig. 4H’, K’) and remained discontinuous in subapical bands in 1.5-fold embryos (Fig. 4H”, K”), suggesting that PKC-3 is required for the correct timing and organization of SJ proteins. In Stage 3 PAR-3^gut(-)^ intestines, DLG-1 also failed to localize (Fig. 4I) and PAR-1 localized weakly to the membrane (Fig. 4L). DLG-1 initially localized as discontinuous patches throughout the intestine of Stage 4 PAR-3^gut(-)^ embryos (Fig. 4I’) and later organized into rings at 1.5-fold (Fig. 4I”). PAR-1 also localized in rings, first primarily in the posterior regions of Stage 4 PAR-3^gut(-)^ intestines (Fig. 4L’) and then throughout the intestine at 1.5-fold (Fig. 4L”). The PAR-1 and DLG-1 structures resembled the HMR-1 and AFD-1 rings described above but assembled at later timepoints (Fig. 4F’ compared with Fig. 4L’), indicating that timely septate junction formation requires PAR-3 and PKC-3, but that some degree of PAR-3 and PKC-3-independent junction assembly occurs in later stages and providing further evidence that the AJ and SJ proteins contribute to different structures.

### PAR-3 and PKC-3 are not required for apical exclusion of LET-413

In addition to roles in junction positioning and maintenance, aPkc phosphorylates basolateral proteins, thereby removing them from the apical domain in many epithelial tissues (Betschinger et al., 2003; Castiglioni et al., 2020; Ventura et al., 2020; Yamanaka et al., 2003). We therefore asked if PKC-3 or PAR-3 is required during basolateral domain establishment in the embryonic intestine. LET-413 membrane localization was similar to controls in Stage 3 PAR-3^gut(-)^ or PKC-3^gut(-)^ embryos (Fig. 5A-C). In Stage 4 and 1.5-fold control embryos, LET-413 was excluded from the apical surface and localized to junctions and basolateral membranes (Fig. 5A’, A’’, (Legouis et al., 2000). As in control embryos, LET-413 was excluded from much of the apical surface of Stage 4 and 1.5-fold PKC-3^gut(-)^ intestines (Fig. 5B’, B’’). Surprisingly, LET-413 was also excluded from parts of the membrane in PAR-3^gut(-)^ embryos. These regions of LET-413 exclusion were initially flanked by DLG-1 puncta (Stage 4, Fig. 5C’ Arrowhead, inset) and eventually surrounded by DLG-1 rings (1.5-fold, Fig. 5C’’). As there is no continuous apical surface in PAR-3^gut(-)^ embryos, we did not observe continuous exclusion of LET-413 along the intestinal midline. In both PAR-3^gut(-)^ and PKC-3^gut(-)^ embryos, the junctional LET-413 enrichment was reduced (Fig. 5B’’, C’’). Together, these results indicate that PAR-3 and PKC-3 are required for the junctional, but not basolateral organization of LET-413 and suggest that localization of PKC-3 by PAR-3 to the apical surface is not required for local basolateral domain establishment.

**Figure 5.**
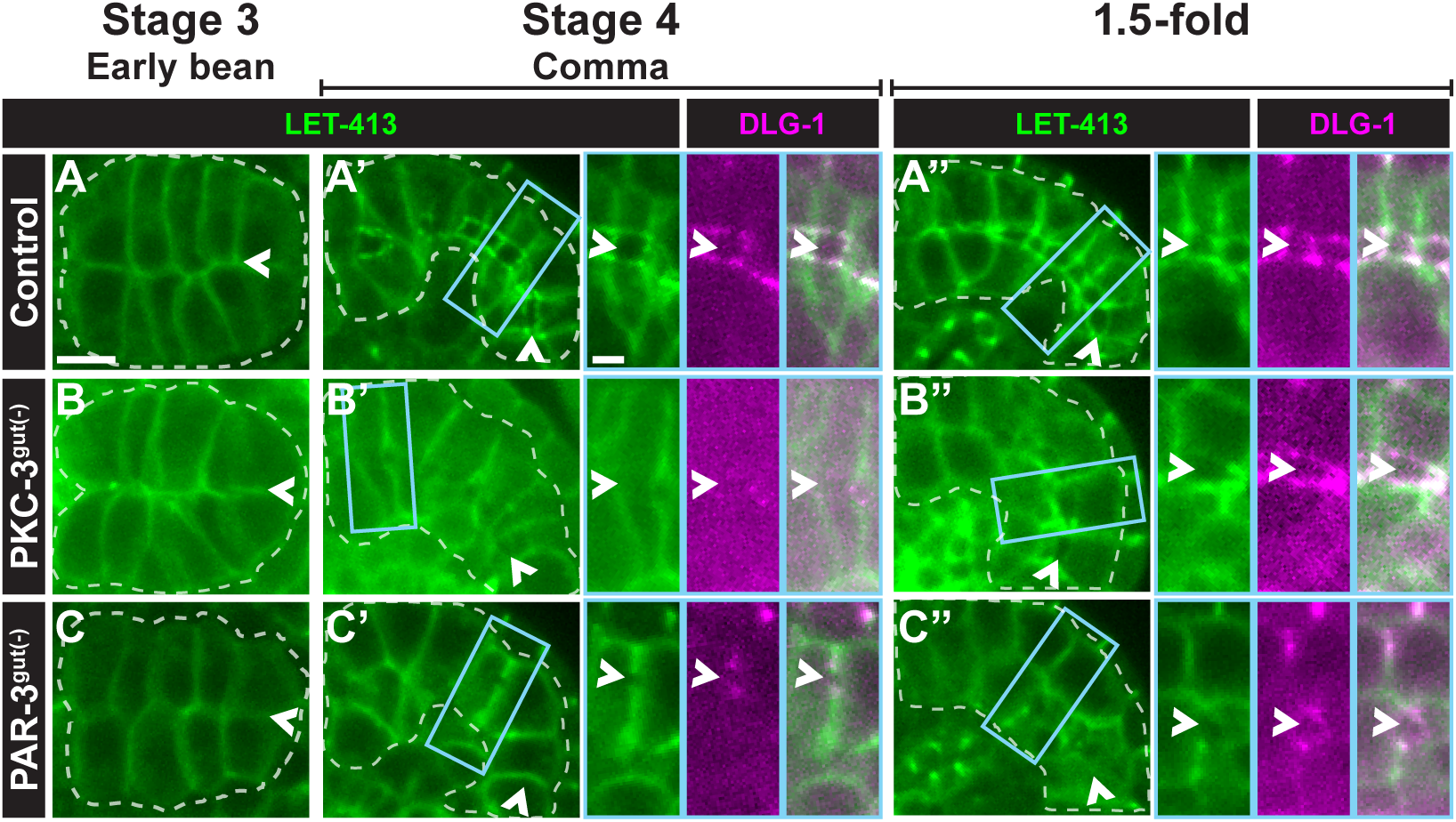
PAR-3 and PKC-3 are not required for apical exclusion of LET-413. Dorsal (Stage 3) and dorsolateral (Stages 4 and 1.5-fold) live images of endogenously tagged LET-413 (green) in control (A-A’’), PAR-3^gut(-)^ (B-B’’), or PKC-3^gut(-)^ (C-C”) embryos. Embryos in A’-C” co-express endogenously tagged DLG-1 (magenta), with magnified view of boxed region shown at right. All images are maximum intensity projections from live imaging. Intestines outlined by white dashed lines, midlines indicated by arrowheads. Scale bar = 5 μm for panels and 2 μm for boxed insets. Stage 3 (Control, n=5; PAR-3^gut(-)^, n=7; PKC-3^gut(-)^, n=8); Stage 4 (Control, n=11; PAR-3^gut(-)^, n=5; PKC-3^gut(-)^, n=11); 1.5-fold (Control, n=11; PAR-3^gut(-)^, n=8; PKC-3^gut(-)^, n=15).

### Aspects of intestinal polarity are established locally but not globally upon PAR-3 depletion

The unexpected assembly of junctional rings in 1.5-fold PAR-3^gut(-)^ embryos raised the possibility that other aspects of apico-basolateral polarity could be established in the absence of PAR-3. To test this hypothesis, we examined apical, junctional, and basolateral structures in PAR-3^gut(-)^ 1.5-fold to 3-fold stage embryos and in newly hatched L1 larvae. In 1.5-fold control embryos, both tubulin and actin localized along the apical surface (Fig. 6A), consistent with the presence of apical microtubules, microvilli, and the terminal web (Carberry et al., 2009; Feldman and Priess, 2012; Leung et al., 1999; MacQueen et al., 2005). In 1.5-fold PAR-3^gut(-)^ intestines, we found discontinuous spheres of actin from which microtubules appeared to radiate (Fig. 6B, (Achilleos et al., 2010; Feldman and Priess, 2012). In control 3-fold embryos and L1 larvae, the actin-binding protein EPS-8, which localizes to the tips of and is required to form microvilli (Croce et al., 2004), localized to the apical surface and was encircled by AFD-1 in the ladder-like junctions (Fig. 6C, E). In 3-fold PAR-3^gut(-)^ embryos and L1 larvae, AFD-1 also formed junctional rings that surrounded large spheres of EPS-8 (Fig. 6D, F). The spheres of EPS-8 and AFD-1 corresponded with the intestinal edemas observed in PAR-3^gut(-)^ L1 larvae in some places (Fig. 3C, data not shown), but were also observed where edemas were not evident. Additionally, LET-413 appeared to be excluded from these apical-like regions, localizing with DLG-1 in PAR-3^gut(-)^ L1 larvae, similar to controls (Fig. 6G, H). Thus, small groups of cells appeared to assemble small, correctly polarized regions--apical domains bounded by junctions and excluding basolateral proteins--in the absence of PAR-3, suggesting an ability to establish local but not global tissue polarity.

**Figure 6.**
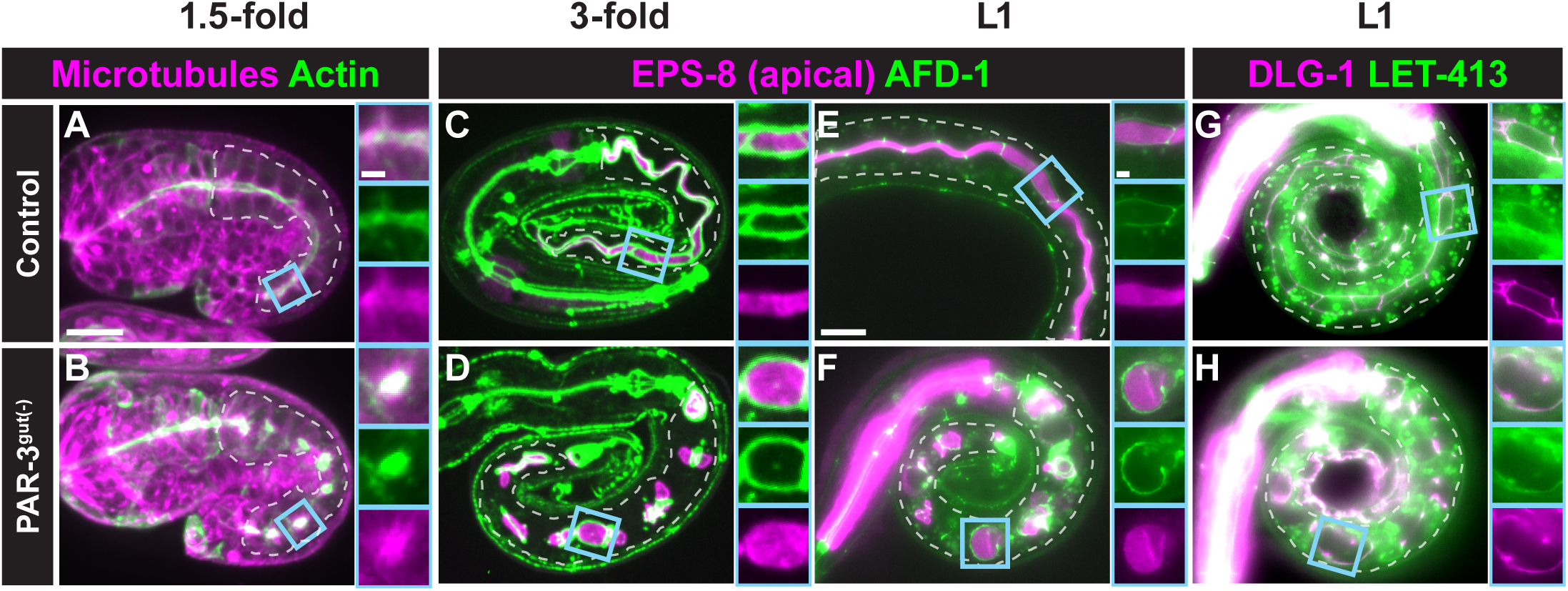
Local apico-basolateral polarity is established in the absence of PAR-3 A-B) Localization of ACT-5/actin (green) and TBA-1/α-tubulin (magenta) transgenes in live 1.5-fold control (n=1) and PAR-3^gut(-)^ (n=10) embryos, using confocal microscopy. C-H) Localization of endogenously tagged indicated proteins in fixed control (n=6) (C) and PAR-3^gut(-)^ (n=7) (D) 3-fold embryos, using confocal microscopy and live control (E, G) and PAR-3^gut(-)^ (F, H) L1 larvae, using a compound microscope. Magnified views of boxed regions are shown at right of each panel. Scale bar = 10 μm for panels and 2 μm for boxed insets. (Control (EPS-8;AFD-1/afadin, n=13; DLG-1;LET-413/Scribble, n=10); PAR-3^gut(-)^ (EPS-8;AFD-1/afadin, n=26; DLG-1;LET-413/Scribble, n=14)

### HMR-1 is required for local polarity establishment in the absence of PAR-3

As junctional rings formed following PAR-3 depletion, we hypothesized that the junctions themselves could establish local cell polarity. We first tested whether HMR-1 is required for the assembly of junctions in the intestine. Both AFD-1 and DLG-1 localized in 1.5-fold HMR-1^gut(-)^ embryos (Fig. 7B), albeit into more broken structures than the ladder-like junctions in controls (Fig. 7A-B). These structures were more continuous than in 1.5-fold PAR-3^gut(-)^ embryos (Fig. 7C). We next co-depleted HMR-1 and PAR-3 in the intestine ([HMR-1;PAR-3]^gut(-)^) to test if HMR-1 is required for the organization of junctions in the absence of PAR-3. We found that junctional rings were completely abolished in 1.5-fold [HMR-1;PAR-3]^gut(-)^ embryos with AFD-1 and DLG-1 puncta randomly localized throughout the intestine (Fig. 7D), in striking contrast to depletion of HMR-1 or PAR-3 alone.

**Figure 7.**
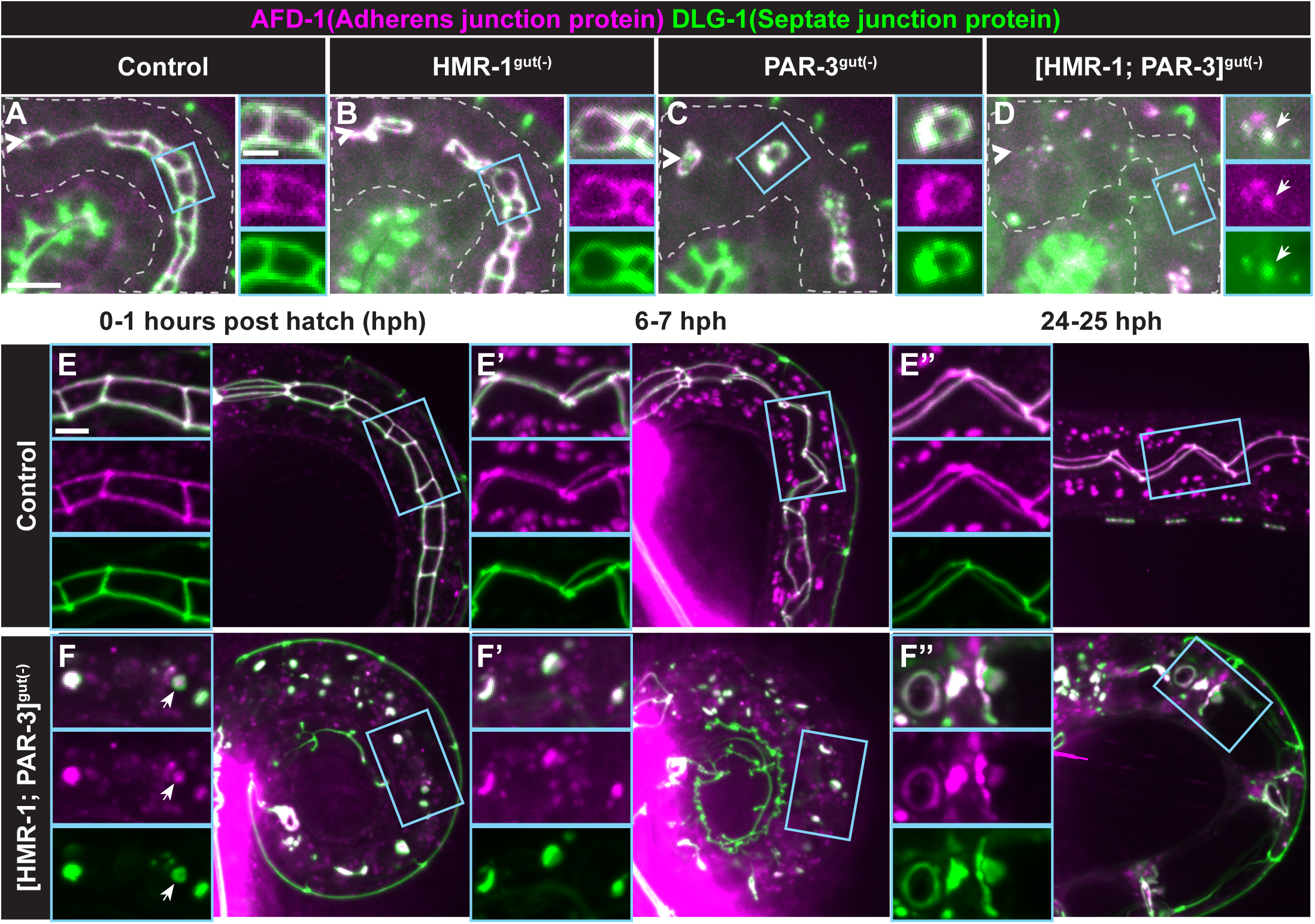
HMR-1 is required for local polarity establishment in the absence of PAR-3. Live images of endogenously tagged AFD-1 (magenta) and DLG-1 (green) localization. A-D) Dorsolateral images of 1.5-fold intestines from control (n=4), HMR-1^gut(-)^ (n=9), PAR-3^gut(-)^ (n=5), or [HMR-1;PAR-3]^gut(-)^ (n=5) embryos. Magnified views of boxed regions are shown at right of each panel. E-F’’) Time courses in control or [HMR-1;PAR-3]^gut(-)^ L1 larvae for hours post-hatching (hph). Magnified views of boxed regions shown at left of each panel. DIC images of [HMR-1;PAR-3]^gut(-)^ included below corresponding fluorescent confocal images in F-F’’. Scale bar = 5 μm for top panels, 10 μm for bottom panels, and 2 μm for boxed insets. (Control (0-1 hph, n=17; 6-7 hph, n=13; 24-25 hph, n=9); [HMR-1;PAR-3]^gut(-)^ (0-1 hph, n=17; 6-7 hph, n=13; 24-25 hph, n=9))

To better understand local polarity in [HMR-1;PAR-3]^gut(-)^ larvae, we followed individual L1 animals over time. In newly hatched (0-1 hours post hatching (hph)) [HMR-1;PAR-3]^gut(-)^ L1 larvae, AFD-1 and DLG-1 appeared largely co-localized in puncta (Fig. 7F). We occasionally observed a hollowing of DLG-1 away from the center of the puncta in these worms. By 6 hph, the same L1 larvae showed an expansion of edemas throughout the intestine, but AFD-1 and DLG-1 remained largely co-localized in puncta and sometimes in small rings that did not surround detectable edemas (Fig. 7F’). Surprisingly, as these L1s aged (24-32 hph), AFD-1 and DLG-1 often formed rings that partially or completely enclosed edemas (Fig. 7F’’). Edemas appeared prior to the formation of junctional rings, suggesting that these rings may arise stochastically through limited space and protein-protein interactions. The asymmetric organization of AFD-1 and DLG-1 first in puncta and then in rings suggests that aspects of local polarity may still arise in the absence of both PAR-3 and HMR-1, which may result from stochastic interactions and highlight the propensity of simple systems to polarize.

## DISCUSSION

Through the live analysis of polarization and tissue specific depletion in the *C. elegans* embryonic intestine, we establish a paradigm where epithelial polarity at the local and global levels are temporally and genetically separable. Local polarity is initiated prior to global polarity before all cells are aligned across a common axis. PAR-3 is required for global polarity establishment across the tissue but is dispensable for local polarity which can still be established through HMR-1-dependent mechanisms. Our results indicate that local polarity establishment is robust but is insufficient to establish global apico-basolateral polarity without additional tissue-level coordination by PAR-3.

### Local vs. global polarity establishment

We found *in vivo* evidence that apico-basolateral polarity is established at two separable levels: 1) locally, individual epithelial cells define an apico-basolateral axis; 2) globally, all epithelial cells within a tissue align their polarity axes in the same orientation, thereby creating a continuous apical surface.

The anatomy of the *C. elegans* intestine allowed us to visualize the natural temporal separation between local and global polarization. Unlike MDCK cysts where cells division across the future apical surface guides local and global apical establishment via the remnant midbody, E8 intestinal cells divide parallel to, instead of across, the future apical surface, requiring a ninety-degree rotation of structures such as the centrosomes and attached nucleus to arrive at the apical surface (Feldman and Priess, 2012; Leung et al., 1999). Additionally, apical proteins are initially positioned as lateral puncta on the membranes of adjacent non-sister cells at E16 and must move to the future apical surface. The asymmetric localization of lateral puncta suggests that local polarization events begin well before global polarity establishment. However, basolateral proteins were not excluded from either the lateral puncta or the nascent apical surface, indicating that although local polarization is an early process, the definition of a basolateral surface normally appears concomitant with global polarity establishment. The separation of local and global polarity may be common among epithelia as the mammalian PAR-3 ortholog similarly localize to asymmetric patches on lateral membranes in the polarizing mouse kidney nephron; these patches are also not initially oriented toward the future apical surface and do not exclude basolateral proteins (Yang et al., 2013).

Local and global polarity are also genetically separable. Degradation of PAR-3 initially impeded local polarity as AJ proteins did not form lateral puncta, and instead localized weakly along lateral cell membranes. However, local polarity was established at later timepoints when HMR-1-dependent junctional rings surrounded proteins associated with apical microvilli and microtubules and excluded basolateral proteins. These locally polarized regions were insufficient to establish global polarity, indicating that PAR-3 is ultimately required to coordinate global but not local polarity establishment. In the embryonic mouse kidney, Afadin appears to similarly be required for global but not local polarity establishment, though Afadin/AFD-1 appears to play a different role in the *C. elegans* intestine as both local and global polarity were eventually established in AFD-1^gut(-)^ embryos ((Yang et al., 2013), Fig. S5).

Separation of local and global polarity may highlight different protein requirements in the evolution of multicellularity and explain the robustness of local polarity. In single cell contexts, local polarity establishment is robust. For example, in *S. cerevisiae*, stochastic changes in protein concentrations at the plasma membrane are sufficient to drive polarity establishment in the absence of other cues (Chiou et al., 2017; Wu and Lew, 2013). In the one-cell *C. elegans* zygote, several redundant processes ensure the robustness of anterior-posterior polarity establishment such that loss of individual polarity proteins or processes (e.g. PAR-2, ECT-2, or LGL-1, actomyosin contractility) does not prevent polarization (Hoege et al., 2010; Motegi and Seydoux, 2013; Motegi et al., 2011; Zonies et al., 2010). However, coordinate polarity orientation of multiple individual cells is less likely to occur correctly through stochastic processes, so global polarity may be more prone to disruption without additional reinforcing mechanisms.

The mechanisms required for early local polarization of lateral puncta are unknown. Unlike in MDCK cysts, midbodies are unlikely to provide a polarizing cue since they are positioned on opposite membranes from lateral puncta (Bai et al., 2020; Li et al., 2014; Lujan et al., 2017). Instead, mitosis from E8 to E16 may promote formation of lateral puncta by trafficking polarity proteins along astral microtubules to their lateral attachment sites or by increasing the concavity of the cell membrane which can promote local accumulation of polarity proteins (Chiou et al., 2017; Feldman and Priess, 2012). Alternatively, as transmembrane proteins at yeast bud scars provide polarizing cues, earlier intestinal cell divisions may locally enrich transmembrane proteins like cadherins, nectins, or claudins at lateral membranes which could then recruit apical and junctional proteins (Chiou et al., 2017). Recently, PAR-3 and other apical and junctional proteins have been found to undergo liquid-liquid phase separation, which is sufficient to promote puncta formation (Liu et al., 2020; Rouaud et al., 2020). Thus, interaction of PAR-3 with transmembrane and other polarity proteins, phase separation, and simple reaction-diffusion may contribute to the establishment of local polarity puncta (Chiou et al., 2017; Wu et al., 2018).

The mechanisms that move lateral puncta, midbodies, and basolateral proteins to the future apical surface are also unknown, but the movement of all of these structures toward the midline is consistent with a global polarization event (Achilleos et al., 2010; Bai et al., 2020; Feldman and Priess, 2012). Actomyosin contractility produces cortical anterior flow during anterior-posterior polarity establishment in the 1-cell *C. elegans* zygote and might be expected to similarly generate apical flow in the intestine (Goehring et al., 2011; Munro et al., 2004). However, acute actin disruption did not disrupt movement of lateral puncta toward the apical surface, suggesting that actomyosin contractility is not required for punctum movement (Feldman and Priess, 2012). By contrast, disruption of microtubules strongly delayed the movement of lateral puncta, suggesting that microtubules are at least partially required for global polarization (Feldman and Priess, 2012).

### PAR-3 and PKC-3 have separable roles in polarity establishment and maintenance

Par3, Par6, and the kinase, aPkc, compose the apical Par complex that is critical for establishing apico-basolateral polarity in many epithelial tissues (Harris and Peifer, 2004; Harris and Peifer, 2005; Hutterer et al., 2004; Totong et al., 2007; Von Stetina and Mango, 2015). While depletion of PAR-3, PAR-6, or PKC-3 from the embryonic intestine resulted in similar L1 larval phenotypes including intestinal obstruction, the absence of a continuous lumen, and edema formation (this study, (Sallee et al., 2021)), we found that these terminal phenotypes arise for distinct reasons. Although, we cannot rule out the possibility that our depletion efficiency varied between proteins, the differences between the PKC-3 and PAR-3 phenotypes are consistent with previous observations (Achilleos et al., 2010; Sallee et al., 2021; Totong et al., 2007).

We found that, although PKC-3^gut(-)^ L1 larvae had severe defects that led to 100% arrest and death, PKC-3 is not required for polarity establishment but for the remodeling and maintenance of global polarity like its binding partner PAR-6 (this study, (Sallee et al., 2021)). These defects may arise during morphogenesis because of a failure to remodel apical junctions, the apical surface, or both. PKC-3 and PAR-6 localize interdependently (Montoyo-Rosario et al., 2020; Sallee et al., 2021; Totong et al., 2007), so correct SJ protein positioning may be achieved by the binding of PKC-3 to the PAR-6 PDZ domain or by PKC-3 directly phosphorylating SJ proteins, similar to the role of aPkc in tight junction organization in mammalian tissues (Elbediwy et al., 2019; Iden et al., 2012). The SJ proteins DLG-1 and PAR-1 appear fragmented both in PKC-3^gut(-)^ embryos and in the *C. elegans* seam cells (this study, (Castiglioni et al., 2020)), suggesting that PKC-3 may be broadly required to organize SJ proteins across epithelial tissues.

In contrast to PKC-3^gut(-)^ embryos, we found a severe delay in local polarization and complete disruption of global polarity establishment following PAR-3 depletion. PAR-3 is a scaffolding protein with an N-terminal oligomerization domain and PDZ domains required for formation of multimeric PAR-3/PAR-6/PKC-3 complexes (Chen and Zhang, 2013; Dickinson et al., 2017). The kinase activity of PKC-3 is thought to be inactive when in complex with PAR-3, and one role of PAR-3 may be to keep PKC-3 in an inactive state (Soriano et al., 2016). Thus, depletion of PAR-3 could allow excess or mislocalized PKC-3 activity, producing a very different phenotype than PKC-3 depletion. Alternatively, the PDZ and C-terminal domains of PAR-3 interact with multiple junctional proteins and membrane lipids, which may allow PAR-3 to position apical and junctional proteins independently of its interaction with PKC-3 localization and activity (Chen and Zhang, 2013). Additionally, PAR-3 may regulate additional kinases such as the Pak1 orthologs MAX-2 or PAK-1 to establish polarity, as in the fly follicular epithelium and cultured mammalian intestinal cells (Aguilar-Aragon et al., 2018). Thus, PAR-3 may coordinate global polarity by positioning apical and junctional proteins as it spreads along the intestinal midline, explaining the more severe defects of PAR-3 depletion than PKC-3 depletion (this study, (Achilleos et al., 2010)).

### Rules that govern polarity establishment: when is PAR-3 required?

PAR-3 plays an essential role in polarity establishment in certain epithelia but is unnecessary in others, raising the question of in what contexts PAR-3 is necessary for polarity establishment (this study, (Achilleos et al., 2010; Chen et al., 2018; Harris and Peifer, 2004; Shahab et al., 2015; Yang et al., 2013)). Notably, PAR-3 only appears to be required during global polarity establishment in the embryonic *C. elegans* intestine as it is dispensable for polarity maintenance and intestinal function in larval and adult worms (Castiglioni et al., 2020). We speculate that the differential requirements for PAR-3 across developmental times and tissues stems from differences in the asymmetric information available to cells prior to polarity establishment. We propose that PAR-3 may be necessary when polarity domains are established for the first time, but not when cells inherit asymmetric information from precursors or are born into already polarized tissues. For example, asymmetries between cell-contacting and contact-free surfaces are sufficient to drive PAR-3-independent radial polarization in early *C. elegans* embryos (Nance, 2014). PAR-3 is also dispensable for polarization in the *C. elegans* epidermis, which has an asymmetric contact-free surface that might coordinate global polarity (Achilleos et al., 2010). The *Drosophila* follicular epithelium and adult midgut do not require PAR-3 and experience asymmetries at future apical and basal surfaces through asymmetric contact with the germline and/or basal ECM which may provide both physical and chemical cues to coordinate global polarity (Chen et al., 2018; Schneider et al., 2006; Shahab et al., 2015; Tanentzapf et al., 2000). Thus, extrinsic informational cues such as chemical signaling gradients, contact with the ECM, or the presence of external contact-free surfaces can guide global polarization independently of PAR-3 (Pickett et al., 2019). The polarizing intestine does not have a contact-free surface, make heterotypic contacts at the future apical surface, or contact basal ECM, perhaps necessitating PAR-3 for global coordination of polarity in the absence of other cues (Nance et al., 2003; Nance and Priess, 2002; Rasmussen et al., 2013). This role is likely in concert with other developmental information as regions of local polarity in PAR-3^gut(-)^ embryos appeared more centrally localized than would be expected by chance. Indeed, PAR-3 functions together with other asymmetric information in both the *Drosophila* blastoderm and the one-cell *C. elegans* zygote when polarity domains are initially established (Dickinson et al., 2017; Harris and Peifer, 2004; Kemphues et al., 1988; Motegi et al., 2011).

The results presented here demonstrate the power of holistic *in vivo* studies, where polarizing tissues encounter their normal external cues, to understand the many mechanisms that drive local and global polarity establishment in different epithelia. Additional systematic studies of polarity pathways are necessary to understand when and how different proteins contribute to the coordination of local and global polarity. Such studies will undoubtedly provide insight into the evolution of multicellularity as well as prove informative to how changes in polarity protein expression contribute differentially to a multitude of epithelial diseases.

## MATERIALS AND METHODS

### *C. elegans* strains and maintenance

*C. elegans* strains were maintained on Nematode Growth Medium (NGM) plates coated with a lawn of *E. coli* OP50 bacteria at 15-20°C as previously described (Sulston and Brenner, 1974). Embryos and larvae used for experiments were collected from 1-to 2-day old adults, with the exception of PAR-3^gut(-)^ and [PAR-3;HMR-1]^gut(-)^ experiments in which 3-day old adults were occasionally used due to the difficulty in obtaining balancer (-) adults. A full list of strains used in this study is available in Table S1.

### CRISPR cloning and editing

New CRISPR alleles were generated using the self-excision cassette (SEC) method as previously described (Dickinson et al., 2015). The pDD162 plasmid was used to deliver Cas9 and sgRNAs modified using the Q5 Site-Directed Mutagenesis Kit (NEB) for sequence specific CRISPR editing of each gene. Homology guided repair templates were generated by Phusion High-Fidelity DNA polymerase (Thermo Scientific) mediated PCR-amplification of appropriate homology arm sequences for N-terminal, C-terminal, or internal-fluorophore tags. Homology arms were then cloned into an SEC backbone plasmid (pJF250, pDD282, or pLC019) with Gibson assembly (NEBuilder HiFi DNA Assembly Master Mix, NEB). A mixture of the modified SEC repair plasmid (25 ng/μl) and modified Cas9/sgRNA plasmid (50 ng/μl) were injected into both gonad arms of 1-day old *zif-1(gk117)* adult hermaphrodites. Injected worms were recovered and treated as previously described to isolate new CRISPR edits and excision of the SEC (Dickinson et al, 2015). Alleles were tagged with ZF::GFP::3XFLAG, GFP::3XFLAG, or mScarlet::3XFLAG. New CRISPR alleles were backcrossed twice with *zif-1(gk117)* worms prior to use in experiments. New worm lines generated through CRISPR editing are listed in Table S1. All sgRNA and homology arm sequences, plasmids, and primers used to generate new CRISPR alleles are available in Table S2.

### Obtaining gut(-) embryos and L1s

All intestine specific depletion strains were maintained with the ZF::GFP allele over an appropriate balancer (See Table S1), with the exception of LGL-1::ZF::mScarlet, which was maintained unbalanced. To obtain gut(-) embryos and L1s, L4 and young adult hermaphrodites lacking the balancer were picked to a fresh plate and maintained at 20°C overnight. The following day, the balancer (-) adults were transferred to 30 μL M9 on a teflon coated slide, washed three times with M9, and incubated in a humidity chamber for 2.5-3 hours at room temperature to obtain gut(-) embryos of the appropriate stages. Worms were cut open to release the gut(-) embryos, which were then imaged (see below) and scored for defects. To obtain gut(-) L1s, the young balancer (-) adults were transferred to a fresh small NGM plate and allowed to lay eggs at 20°C for 1-4 hours. Adults were then removed and plates returned to 20°C for 12-24 hours when L1s were scored for defects. This strategy depletes both maternal and zygotic proteins from the intestine, as both supplies of the protein is ZF tagged and subject to degradation. Depletion was verified by loss of GFP fluorescence at the intestinal midline (Fig. S3). While the ZF degradation system does not create a null line, it has been used to robustly deplete ZF-tagged proteins (Abrams and Nance, 2021; Liang et al., 2020; Magescas et al., 2021; Sallee et al., 2021; Sallee et al., 2018; Sanchez et al., 2021) and we see strong depletion.

### Immunofluorescence

1-to 2-day old hermaphrodites were incubated in 30 μL M9 in a humidity chamber at room temperature for 2.5-3.5 hours, and were then cut open to release their embryos which were stained as previously described (Leung et al., 1999). Embryos were attached to a poly-L-lysine coated slide with Teflon spacers and rapidly frozen on dry ice. Embryos were permeabilized with the freeze-crack method and fixed in -20°C 100% methanol for 5 minutes. Slides were then washed twice in 1X PBS for 5 minutes at room temperature followed by a single 5-minute wash in 1XPBS with 0.1% Tween (PBT). Embryos were incubated in primary antibody solutions in a humidity chamber at 4°C overnight. Slides were washed three times in PBT for five minutes each, then incubated in secondary antibody solutions in a humidity chamber at 37°C for 1 hour. Slides were washed twice in PBT for five minutes each, followed by a five-minute wash in 1X PBS at room temperature. Vectashield (Vector Laboratories) was added to each slide and coverslips sealed to slides with fingernail polish. Slides were imaged on a Nikon Ti-E inverted spinning disk confocal microscope (see below). The following primary antibodies were used: anti-GFP(Abcam (AB6556), 1:200), anti-PAR-3 (DSHB (P4A1), 1:25). The following secondary antibodies were used: Cy3 anti-mouse (Jackson (115165166), 1:200), 488-goat-anti-rabbit (Jackson (111545144) 1:200), 488-goat-anti-mouse (Jackson (115545166), 1:200), 647-donky-anti-rabbit (Jackson (711605152), 1:50). DAPI (Sigma (D9542), 1:10,000) staining was used to visualize nuclei.

### Microscopy

Embryos were collected from 1-to 2-day old hermaphrodites, incubated in M9 for 2.5-3.5 hours at room temperature and used for live imaging. For [PAR-3;HMR-1]^gut(-)^, embryos were picked off of small plates containing adult hermaphrodites that lacked the hT2 balancer. Embryos were imaged live and mounted on 3% agarose pads in 1X M9. Images were acquired on a Nikon Ti-E inverted microscope (Nikon Instruments, Melville, NY) with a 60X Oil Plan Apochromate (NA = 1.4), an Andor Ixon Ultra back thinned EM-CCD camera, using 405 nm, 488 nm, and/or 561 nm lasers, and a Yokogawa X1 confocal spinning disk head objective controlled by NIS Elements software (Nikon). Images were acquired at a z-sampling rate of 0.3 – 0.5 μm. Some L1 larvae were imaged using a Nikon Ni-E compound microscope with a 100X Oil Plan Apochromat (NA = 1.45) objective. Maximum intensity projections were made with FIJI. Imaris was used for 3D rotation.

### Image Analysis and Statistical Analysis

#### Quantification of midline enrichment and basolateral exclusion

To determine if basolateral proteins were enriched at the midline and when they became excluded from this surface, we analyzed images from live embryos at Stage 1, Stage 2, Stage 2.5/3, and Stage 4. Using FIJI, we used endogenous PAR-3::mCherry or *end-1*p-driven BFP::CAAX signal to select the brightest plane to define the intestinal midline and made a sum Z-projection of 3 slices around this plane for further analysis. 2-μm wide boxes were drawn by hand at the apical and lateral surfaces to define ROIs for basolateral proteins. Apical enrichment was determined by dividing the average apical signal by the average lateral signal for each embryo. A 1-μm wide line segment was drawn across the midline at Int2 and pixels within 5-μm of the midline were selected to generate a line profile. Plots of apical enrichment and profile plots were generated and normalized in R with ggplot2.

#### Quantification of distance between junctional intensity peaks

We used FIJI to determine the distance between the left and right sides of the junctions across the midline. For comparisons of AJ proteins and SJ proteins, we analyzed 1.5-fold embryos expressing two endogenously tagged junctional proteins, each tagged with a different fluorophore. A sum Z-projection of 5 slices was made around the brightest intestinal plane based on the GFP or mNeonGreen junctional signal. Three 1-μm wide line segments were drawn across the midline between Int5 and Int8 to generate line profiles for both junctional channels. Measurements were made near the middle of each intestinal ring in order to accurately measure the distance between the left and right sides of the junction while avoiding junctional signal between anterior-posterior neighbors. The difference between the two brightest peaks (highest signal intensity) for each line profile was determined to find the distance between left and right sides. The distances were averaged from the three profile lines for each embryo. GraphPad was used to conduct paired two-tailed t-tests to determine if there were statistically significant differences in the distances between the paired junctional components (HMR-1 vs DLG-1, HMR-1 vs AFD-1, AFD-1 vs DLG-1, AFD-1 vs PAR-1, DLG-1 vs PAR-1).

### L1 Arrest and Larval Growth Assays

To assess embryonic lethality and larval growth, 10 L4 hermaphrodites lacking the balancer and 30 control L4 hermaphrodites (10 with the ZF::GFP tag alone, 10 *zif-1(gk117)*, and 10 *zif-1(gk117);*(Intestine specific ZIF-1)) were picked to fresh plates and maintained at 20°C overnight. The following day, 1-day old adults were singled onto 10 plates and allowed to lay embryos for 4 hours at 20°C. Adult worms were then removed and embryos were counted at least twice. Plates were returned to 20°C for 24 hours. Embryos were counted again and scored as unhatched/dead. The difference between embryos counted on the first day and embryos counted 24 hours later was used to determine the number of L1 larvae on each plate. Plates were returned to 20°C for 24 hours. Embryos, L2 larvae, and L3 larvae were counted at least twice and then plates were returned to 20°C for 24 hours. On the final day (72 hours after adult hermaphrodites were singled), all larval stages (L1-adult) were counted. After 72 hours most L1 larvae were still alive but appeared sick, only moving upon touch stimulation.

### L1 SMURF Feeding Assays

1-to 2-day old adult hermaphrodites were picked to fresh, small NGM plates and allowed to lay embryos for up to 16 hours at 20°C. Hatched L1 larvae were picked into a 30 μL drop of standard overnight OP50 bacterial culture and 10μL of 20% blue food coloring (FD and C Blue #1 Powder, Brand: FLAVORSandCOLOR.com, Amazon) in water was added to the bacteria. L1s were incubated in the blue dyed OP50 in a humidity chamber at room temperature for 3 hours. To collect larvae, the dyed OP50 solution was transferred and spread over a medium NGM plate. L1s were located using oblique illumination and were then mounted in 2 mM levamisole on a 3% agarose pad and imaged on a Nikon Ni-E compound microscope with a 100X Oil Plan Apochromat (NA = 1.45) objective and an additional (2X) digital zoom with a Google Pixel 4, mounted to the microscope eyepiece. Rarely, an L1 was damaged in the process of transferring it to levamisole. These larvae appeared desiccated, and their entire bodies were dyed blue. Damaged L1s were excluded from analysis.

## DATA AVAILABILITY

All data generated or analyzed during this study are included in the manuscript.

## ACKNOWLEDGEMENTS

We thank Jeremy Nance, Mike Boxem, Renaud Legouis, Ken Kemphues, Dan Dickinson, and Lauren Cote for worm strains and plasmids. We also thank members of the Feldman lab for helpful discussions about the research and manuscript. Some strains were provided by the CGC, funded by the NIH Office of Research Infrastructure Programs (P40 OD010440). This work was funded by F32 GM129900-01 awarded to MAP, F32 GM120913-02 and K99 GM13548901 to MDS, T32 GM007276 and a Stanford Graduate Fellowship to VFN, and an NIH New Innovator Award DP2 GM119136-01 and RO1 GM133950 awarded to JLF. KS is an investigator in the Howard Hughes Medical Institute,

